# Genome Report: A blue mussel chromosome-scale genome assembly for aquaculture, marine ecology and evolution

**DOI:** 10.1101/2022.11.17.516937

**Authors:** Tim Regan, Tiago S. Hori, Tim P. Bean

## Abstract

The blue mussel, *Mytilus edulis* is part of the *Mytilus edulis* species complex, encompassing at least three putative species: *M. edulis, M. galloprovincialis* and *M. trossulus*. These three species occur on both sides of the Atlantic and hybridize in nature, and both *M. edulis* and *M. galloprovincialis* are important aquaculture species. They are also invasive species in many parts of the world. Here, we present a chromosome-level assembly of *Mytilus edulis*. We used a combination of PacBio sequencing and Dovetail’s Omni-C technology to generate an assembly with 14 long scaffolds containing 94% of the predicted length of the *M. edulis* genome (1.6 out of 1.7 Gb). Assembly statistics were total length 1.65 Gb, N50 = 116 Mb, L50 = 7 and, L90 = 13. BUSCO analysis showed 92.55% eukaryote BUSCOs identified. AB-*Initio* annotation using RNA-seq from mantle, gills, muscle and foot predicted 47,128 genes. These gene models were combined with Isoseq validation resulting in 65,505 gene models and 129,708 isoforms. Using GBS and shotgun sequencing, we also sequenced 3 North American populations of *Mytilus* to characterize single-nucleotide as well as structural variance. This high-quality genome for *M. edulis* provides a platform to develop tools that can be used in breeding, molecular ecology and evolution to address questions of both commercial and environmental perspectives.

## Introduction

The blue mussel (*Mytilus edulis*) is common to the North Atlantic from Arctic to Mediterranean regions, with habitat ranging from upper shore to the shallow subtidal (Hayward and Ryland 2017). *M. edulis* is known to hybridise with *M. trossulus* in North America and with *M. trossulus* and *M. galloprovincialis* in Europe. Together, these three species form the *M. edulis* species complex (**Fig. 1**) (McDonald et al. 1991).

**Fig. 1.**
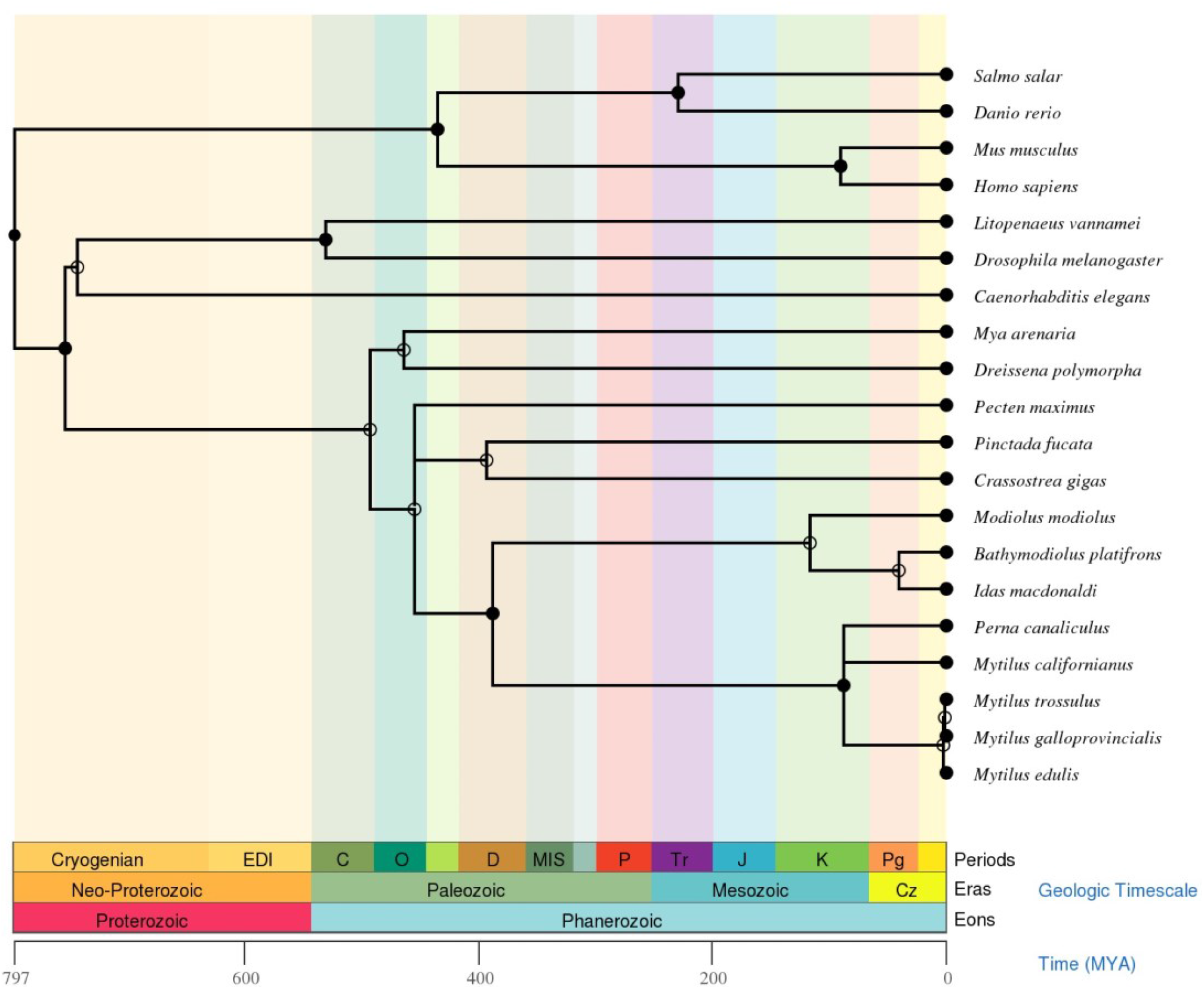
TimeTree for *Mytilus edulis*. Mytilidae diverged ∼387 MYA. Generated using TimeTree (Kumar et al. 2022).

This reef-building bivalve is an ecosystem engineer. Blue mussels dominate fouling communities in shallow and substrata providing important secondary habitat (Norling and Kautsky 2007). Offshore wind energy structure surveys found that they can cover the structures with up to 3.4 kg of biomass m^−2^ (Krone et al. 2013). Through filter-feeding, eutrophication is reduced which can alter ecosystems (Broszeit et al. 2016). This nutrient cycling ability has been harnessed by using *M. edulis* to study the fate of persistent organic and metal pollutants (Chase et al. 2001; McEneff et al. 2014), for the bioremediation of waste (Broszeit et al. 2016) and reduce environmental effects from salmon farms (MacDonald et al. 2011).

Mussels, a key bivalve production species (FAO 2020), face decreasing wild spat availability for aquaculture in the UK and elsewhere (Regan et al. 2021). These losses are attributed to multiple stressors including warmer seas (Seuront et al. 2019) causing a poleward range contraction (Jones et al. 2010). Additionally, warmer oceans elevate dissolved CO_2_ leading to Ocean Acidification (OA) impacting mussel viability (Asplund et al. 2014) and disease resistance (Ellis et al. 2015). This makes mussels more susceptible to bacterial pathogenesis (Eggermont et al. 2017; Ripabelli et al. 1999), with emerging pathogens posing a constant threat (Charles et al. 2020; Cano et al. 2022). Infectious disease such as disseminated Neoplasia (DN) of *M. trossulus* origin is associated with reduced fitness in *Mytilus spp*. (Burioli et al. 2021). Furthermore, the effects of hybridisation between the three species of the *Mytilus edulis* complex are yet uncertain with suggested negative effect of *M. trossulus* hybridisation and potential adaptive introgression in the case of *M. galloprovincialis* hybridisation (Fraïsse et al. 2014; Kenchington et al. 2020; Michalek et al. 2021).

To protect the aquaculture industry from these threats, hatchery efforts have been launched in the UK (Regan et al. 2021), elsewhere in Europe (Kamermans et al. 2013) and in Canada (Gurney-Smith et al. 2017). However, these efforts have not been straightforward and a better understanding of fundamental biology is required to achieve commercial success. Despite their importance in aquaculture and the valuable ecosystem services they provide, no chromosomal assembly existed for any species within the *Mytilus edulis* species complex prior to this study. Improved genomic tools are required to address fundamental biological questions such as inheritance patterns and adaptations.

Like many bivalves, the mussel genome is highly heterozygous (3.5%) with an estimated 43% repeat content. The linear plot of k-mer abundance analysis clearly shows a heterozygous peak in addition to the homozygous peak and estimates a shorter haploid genome length of 1.18 Gb compared to flow-cytometry data. The estimated repeat content of the *M. edulis* genome is ∼43%. These characteristics make assembly of these genomes challenging. However, recently genomes for the American oyster, Pacific Oyster, *M. corruscus* and *M. califoniaus* have been assembled to chromosome-level using short and long-read sequencing technologies as well as Hi-C-based scaffolding (Yang et al. 2021; Paggeot et al. 2022a; Penaloza et al. 2021; Gomez-Chiarri et al. 2015). We used a similar approach in this project to produce a highly contiguous assembly. Practical application of this assembly is demonstrated in cross-species synteny analyses and in population structure of *Mytilus* individuals sampled from different regions of the Canadian Atlantic.

## Methods

### DNA extraction, library preparation for genome assembly

One naïve blue mussel sample (Anne) was selected from samples collected by the Provincial Department of Communities and Fisheries in the estuary Foxley river in PE. Foxley River was selected as a sampling site because there is no grow-out aquaculture there (i.e. seed from other bays are not transferred to the area). Due to potential introgression of other species of the *Mytlius* species complex (e.g. *M. trossulus*), this sample was genotyped using 12 SNPs described by (Wilson et al. 2018). We sampled the gill, mantle and muscle of sample “Anne” aseptically, flash froze fresh tissues in liquid nitrogen and preserved them at −80°C. Tissues were shipped to Dovetail Genomics in Scott’s Valley, CA, in excess dry ice. Dovetail extracted high molecular weight (HMW) DNA using an in-house modified CTAB method. Dovetail prepared and sequenced PacBIO SMRTbell libraries to a depth of 196X using the Sequel II sequencer. Processed PacBio data (from Dovetail) can be found on SRA under accession number SRX11246493.

### Raw contig assembly, scaffold formation and polishing

Dovetail generated a primary contig-level assembly using wtdbg2 (Ruan and Li 2020). The contig-level assembly was filtered of putative duplicated haplotypes and contaminants using Purge Haplotigs (Roach et al. 2018) and Blobtools2 (Challis et al. 2020), respectively. Scaffolding was performed using Omni-C libraries and the HiRise assembler (v1.0). The same DNA sample used by Dovetail was shipped to UWM (University of Wisconsin – Madison) and sequenced to a 100X using the NovaSeq sequencer. These data were trimmed using Trimmomatic (v) and used for polishing with racon (v1.4.3) (Vaser et al. 2017). Completeness was evaluated using compleasm v0.2.5 (Huang and Li 2023) (eukaryota_odb10: 255 BUSCOS, metazoa_odb10: 954 BUSCOS and mollusca_odb10: 5295 BUSCOS) (Manni et al. 2021) and Merqury (Rhie et al. 2020). Merqury analysis was carried out with the same read set used for polishing as the original PacBio CLR reads were not suitable for this analysis. General length metrics were obtained using QUAST (v5.0.1) (Gurevich et al. 2013; Mikheenko et al. 2018). Synteny mapping between the *M. edulis* and the *M. coruscus* (GCA_017311375.1) (Yang et al. 2021) was done using the MCScanX.h function of MCScanX (v2) (Wang et al. 2012). Putative orthologous groups were identified with Orthofinder (v.2.5.4) using predicted gene structures for *M. edulis* (this work) and *M. coruscus* (Yang et al. 2021). Dot plots and circle plots were generated using MCScanX (v.2).

### RNA preparation and Isoseq analysis

Isoseq3 analysis of CSS data from muscle, gill, hemolymph, and foot (2.9 million reads −188,165 Mb) identified 216,343 high-quality putative full-length (FL) transcripts (from 2.8 million reads containing poly-A tails). We shipped flash frozen gill and adductor muscle tissue from sample “Anne” to the Biotechnology Centre Core facility at the University of Wisconsin, Madison (UWM). Samples were homogenized using a Qiagen Tissuelyser (2 min @ 20 Hz). RNA was extracted using the RNeasy Mini Kit (Qiagen) with on-column DNAse treatment. UWM performed RNA QC with a nanodrop and bioanalyzer and prepared libraries using the Iso-seq Library SMRTbell express template prep kit. Libraries were sequenced using one Sequel II SMRT cell in CSS mode (i.e. HiFi reads). UWM provided de-multiplexed processed HiFi reads. RNA samples from foot, mantle and gut were sent to the Genome Excellence Centre (Genome Quebec) in Montreal. HiFi sequencing was carried out as above. Putative full-length transcripts were identified using the IsoSeq3 pipeline (https://github.com/ylipacbio/IsoSeq3). Putative open read frames (ORF) were identified using TransDecoder (v5.5.0) (https://github.com/TransDecoder/TransDecoder/wiki).

### Annotation

Repeat modeller (4.1.0) (Flynn et al. 2020) was used to predict repeat motifs for *M. edulis* and Repeat Masker (2.0.2a) (Smit et al. 2015) was used to mask the final assembly. Ab-initio annotation was done using Augustus (v3.4.0) (Stanke et al. 2006) trained with genes from *Mytilus galloprovincialis, Mytilus coruscus* and *Crassostrea virginica*. Additional hints were generated using short-read RNA-seq (data not shown) and Isoseq data generated herein. The ab-initio annotation was updated using PASA (v2.5) with alignments of a de-novo transcriptome produced using Trinity (v2.8.15) (Grabherr et al. 2011) in Genome-Guided mode. Two runs through PASA were used to update the *ab-initio* annotation. Full-length (FL) transcripts from Isoseq3 pipeline were mapped to the genome using pbmm2 with pre-sets for Isoseq and filtered based on quality Isoseq3 collapse (minimum alignment identity/coverage: 0.90/0.90) and Isoseq3 refine. We used the resulting gff annotation to run SQANTI3 (v4.3.0). We used sqanti_qc.py to generate quality data for sqanti_filter.py. Filtering to remove artifacts was carried out using the default parameters. Lastly, ab-initio predications and filtered Isoseq FLs were merged with AGAT (v.1.2.0). Using agat_sp_merge.pl, we removed duplicate gene models/isoform and assigned orphan isoforms from the Isoseq data when possible. Amino acid sequences were translated from the CDS using agat_sp_extract_sequences using options -t “CDS” -p --cfs --acs –asc.

### Sample collection for SNP discovery and population structure

We collected gill samples for DNA extractions using standard molecular biology techniques and preserved them in non-denatured 100% ethanol. For population genetics analysis, we sampled mussels in sets of 96 from west to east PEI in Foxley River (FOX) (wild population), Malpeque Bay (MB) (North Shore, Prince County); French River (FR), Stanley Bridge (ST), Whetley River (WR), Tracadie (TRC) (North Shore, Queen county); Orwell (OR) (South Shore, Queen’s county; Morell (MRL) (North Shore – King’s county) and; Murray River (MR) (South Shore, King county) (**Fig. 2**). Seed deployed on these sites were originally collected in St. Peter’s bay, Brudenell River, and Malpeque bay. We also collected samples in the Bras D’or Lake in Cape Breton (Nova Scotia), the Magdalene Island (Québec) and Notre Dame bay in Newfoundland.

**Figure 2:**
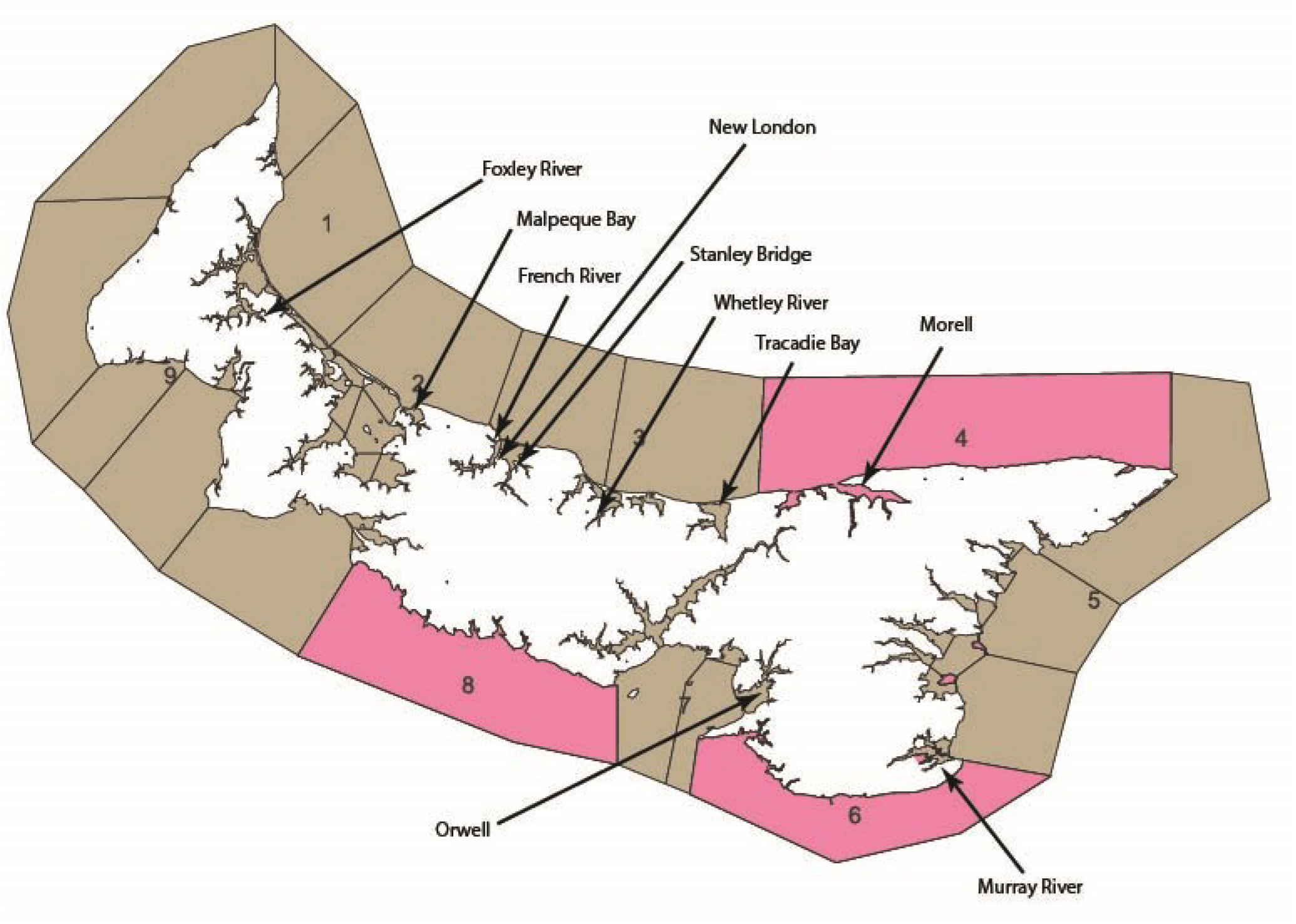
Sampling Locations in PEI.

### SNP markers, Admixture analysis, population structures

Samples for SNP discovery and population genetics were shipped to LGC genomics in Berlin. Restriction-associated DNA libraries (RAD) libraries were prepared by LGC with using MslI, normalized and sequenced using a NextSeq sequencer. The resulting reads were trimmed and checked for the restriction site by LGC. This final read set was used to identify SNPs and call individual genotypes using Tassel5 (v.5.2.4). SNPs were filtered using MAF (<0.01) and percent of individuals with genotypes. For the SNP discovery, the samples from Cape Breton were excluded from filtering analysis. These samples were shown to be a pure *M. trossulus* population and had missing calls for a significant number of sites. For population genetics analysis, a second set of SNPs that were successfully called across all populations was used. All samples were also genotyped for the 12 species discrimination SNPs from (Wilson et al. 2018). All population genetics analyses were performed using the dpca function of the R package Adegenet (v.2.1.7) and STRUCTURE (v.2.3.4) or fastSTRUCTURE (v1.0) (K 3 to K10, 5000 repetitions, 1000 burn in). Analysis of species discrimination SNPs included the genotypes published by (Corrochano-Fraile et al. 2022) as outgroups.

## Results and Discussion

### Genome Assembly and Annotation

The chromosome-level assembly presented herein was produced in two stages. First, ∼15 million PacBio CLR reads (∼340 Gb) were produced, representing coverage of 196X for an estimated genome size of 1.7 Gb (Hinegardner 1974; Rodríguez-Juíz et al. 1996). These reads were assembled into contigs using wtdbg2, which uses uncorrected reads (Ruan and Li 2020). The primary assembly was 1.96 Gb long in 17,825 contigs and a N50 of 443 Kb. After haplotype purging and contaminant removal, the final contig assembly had 10,111 contigs, for a total of 1,65 Gb and a N50 of 518 Kb. Following scaffolding using Omni-C libraries and the HiRise assembler, we generated a primary chromosome-level assembly made of 2,117 contigs. We removed putative contaminants using Blobtools by eliminating sequences coming from non-molluscan organisms. This assembly was further filtered to contain only sequences > 5,000 bp. The resulting draft is deposited on NCBI assembly under accession number GCA_019925275.1. The final assembly is made of 1,119 contigs and has an N50 of 116 Mb. The 14 putative *M. edulis* chromosomes are deposited under accession numbers CM034349.1 to CM34362.1. Detailed statistics for the assemblies can be found in **Table 1**.

**Table 1.** Summary statistics for *M. edulis* assemblies.

|  | Anne (This study) | Corrochano-Fraile *et al.*<br>(2022) | xbMytEdul2.1, Darwin<br>Tree of Life (2024) |
| --- | --- | --- | --- |
| *Length* | 1,659,567,081 bp | 1,827,085,763 | 1,374,471,240 |
| *# of scaffolds* | 1,119 | 3,339 | 2,563 |
| *# of contigs* | 9,866 | 5,966 | 3,754 |
| *N50* | 116,503,180 | 1,097,279 | 1,734,586 |
| *Coverage* | 196.0x | 152.0x | 30.0x |
| *Completeness*<br>*(Merqury)* | 76.71 | NA | NA |
| *QV* | 32.39 | NA | NA |

Compared to the assembly published in Corrochano-Fraile et al. (2022), the assembly presented herein has better contiguity (**Table 1**). Our assembly is also shorter than the assembly from Corrochano-Fraile et al. (2022). The two assemblies’ length falls close to the estimated size of the genome based on c-value (1.7) (Rodríguez-Juíz et al. 1996) and is significantly longer than what is estimated by k-mer abundance analysis with GenomeScope (1.18 Gb).

Despite the total assembly length being close to that estimated using c-values, k-mer-based completeness analysis recovers only 76% in a set of Illumina reads origination from sample “Anne”. When the putative purged haplotigs were added back to the assembly, recovery was ∼83%. This apparent low k-mer recovery is probably a combination of the consensus being different from either haploid genomes, the error rate in the original PacBio data, the fact that the reads used for polishing were not used from the primary assembly, the contigs removed based on length or contaminant status, and the gaps arising from the Omni-C scaffolding. It is also possible that the high heterozygosity affects the accuracy of k-mer abundance analysis, as shown by the large discrepancy between genome size estimates.

Completeness analysis in Merqury resulted in 76.71% recovery of k-mers present in the polishing Illumina data from the final version of the assembly. We also evaluated the completeness of the assembly when combined with purged haplotypes, which was 83.44%. QV value for the primary assembly was 32.39, while the combined draft had a QV of 30.73. Compleasm BUSCO analysis showed a recovery of 92.55%, 91.95%, and 88.5% complete BUSCOS against the eukaryote, metazoan and molluscan databases, respectively. The 14 putative chromosomes represent ∼96% of the assembly, with lengths varying from 140 Mbp to 90 Mbp. These data and the N50 metric show that this assembly has high contiguity and that this assembly and its annotation will be highly useful for aquaculture, evolution and molecular ecology studies. Herein, we illustrate the possible applications of this assembly by performing population and synteny analyses.

### SNP discovery and population structure

Due to the close relationship between the members of the *Mytilus* species complex, we wanted to verify that the individual sampled (Anne) was pure *M. edulis*. Population Structure analysis conducted using 12 SNPs (Wilson et al. 2018) clearly separated *M. edulis* populations in PEI from other regions of Canada. We generated two sets of SNPs: the first set totalling 71,231 SNPs using only samples from PEI and made a polymorphic collection of SNPs in *M. edulis*. The second set, with ∼6,000 markers, is a polymorphic set of SNPs in both *M. edulis* and *M. trossulus*. Population structure and putative admixture are shown in **Figure 4**. In the DPCA and PCA, untrained clustering clearly separated both CB from PEI/NL/MAG and also the putative populations in the Gulf of St. Lawrence. In green, are the majority of samples from NL and MAG, while blue represents individuals from PEI.

We used *M. trossulus, M. galloprovincialis* and European *M. edulis* genotypes as outgroups. The later inclusion of 96 individuals from Cape Breton (NS) showed no significant evidence of *M. trossulus* introgression in PEI samples. Given that Cape Breton has long been considered a pure *M. trossulus* population (Wilson et al., 2018), we are confident that the sample Anne represents an *M. edulis* individual. We also genotyped over 500 PEI individuals and 96 samples from Magdalene Island and 96 individuals from Newfoundland with the same 12 SNP panel. As before no significant introgression of *M. trossulus* was detected in PEI. However, we only genotyped 96 samples collected in an area with no grow-out leases (Foxley River). Although unlikely, we cannot rule out the possibility of a sampling bias in aquaculture sites favouring *M. edulis*. DPCA and Structure analysis indicate that there is low population stratification between different regions of PEI, while the Magdalene islands and NL are distinct populations. The populations from NL and the Magdalene Islands are more similar to each other than from the populations from PEI. For animals from one sampling event at S.t Peter’s bay, we found evidence of shared genetics between PEI and the population to the northeast of the island.

### Annotation and synteny analysis

We identified 196,111 putative open reading frames from the 216,343 FL transcripts using Isoseq3 analysis of CSS data from muscle and gill (2.9 million reads −188,165 Mb). BLASTp analysis against the uniref90 database returned informative hits for ∼80% (164,969) of these translated transcripts. *Ab-initio* gene prediction in Augustus detected 46,604 gene models that produced 46,604 transcripts after filtering based on evidence support. After two rounds of PASA updates, the final number of gene models was 47,128 and the number of transcripts was 55,138. After Isoseq refine and collapse 85,099 isoforms survived and following SQANTI3 filtering 70,592 isoforms remained. They were assigned to 31,211 unique genes. Finally, the combined annotation has 65,505 gene models and 129,708 isoforms. Proteins were translated from the CDS ensuring only complete CDS were translated and that isoforms were not incorrectly fused together. This resulted in 45,379 amino acid sequences. Compleasm BUSCO analysis of these 45,379 proteins showed a recovery of 78.43% and 76.83% complete BUSCOS against the eukaryote and metazoan databases, respectively.

Synteny analysis showed a high degree of collinearity between putative chromosomes of *M. edulis* and *M. coruscus* (**Fig. 3**). However, putative inversions, transposition and deletion can be observed in almost all chromosomes. Gene order in chromosomes represented by sequences CM034349.1 (*M. edulis* chromosome 1) - CM029595.1 (*M. coruscus* LG1) and CM034343.1 (*M. edulis* chromosome 4) – CM029599.1 (*M. coruscus* LG5) showed the highest conservation. Putative orthologous relationships between *M. edulis* chromosomes and *M. coruscus* linkage groups (LG) are shown in **Table 2**.

**Table 2:** Putative synteny between *M. edulis* and *M. coruscus*. Ids are NCBI Assembly database Molecule name and accession number.

| *M. edulis* | *M. coruscus* |
| --- | --- |
| Chromosome 1 (CM034349.1) | LG01 (CM029595.1) |
| Chromosome 2 (CM034350.1) | LG04 (CM029598.1) |
| Chromosome 3 (CM034351.1) | LG03 (CM029597.1) |
| Chromosome 4 (CM034352.1) | LG05 (CM029599.1) |
| Chromosome 5 (CM034353.1) | LG10 (CM029604.1) |
| Chromosome 6 (CM034354.1) | LG06 (CM029600.1) |
| Chromosome 7 (CM034355.1) | LG02 (CM029596.1) |
| Chromosome 8 (CM034356.1) | LG11 (CM029605.1) |
| Chromosome 9 (CM034357.1) | LG07 (CM029601.1) |
| Chromosome 10 (CM034358.1) | LG08 (CM029602.1) |
| Chromosome 11 (CM034359.1) | LG09 (CM029603.1) |
| Chromosome 12 (CM034360.1) | LG14 (CM029608.1) |
| Chromosome 13 (CM034361.1) | LG12 (CM029606.1) |
| Chromosome 14 (CM034362.1) | LG13 (CM029607.1) |

**Figure 3.**
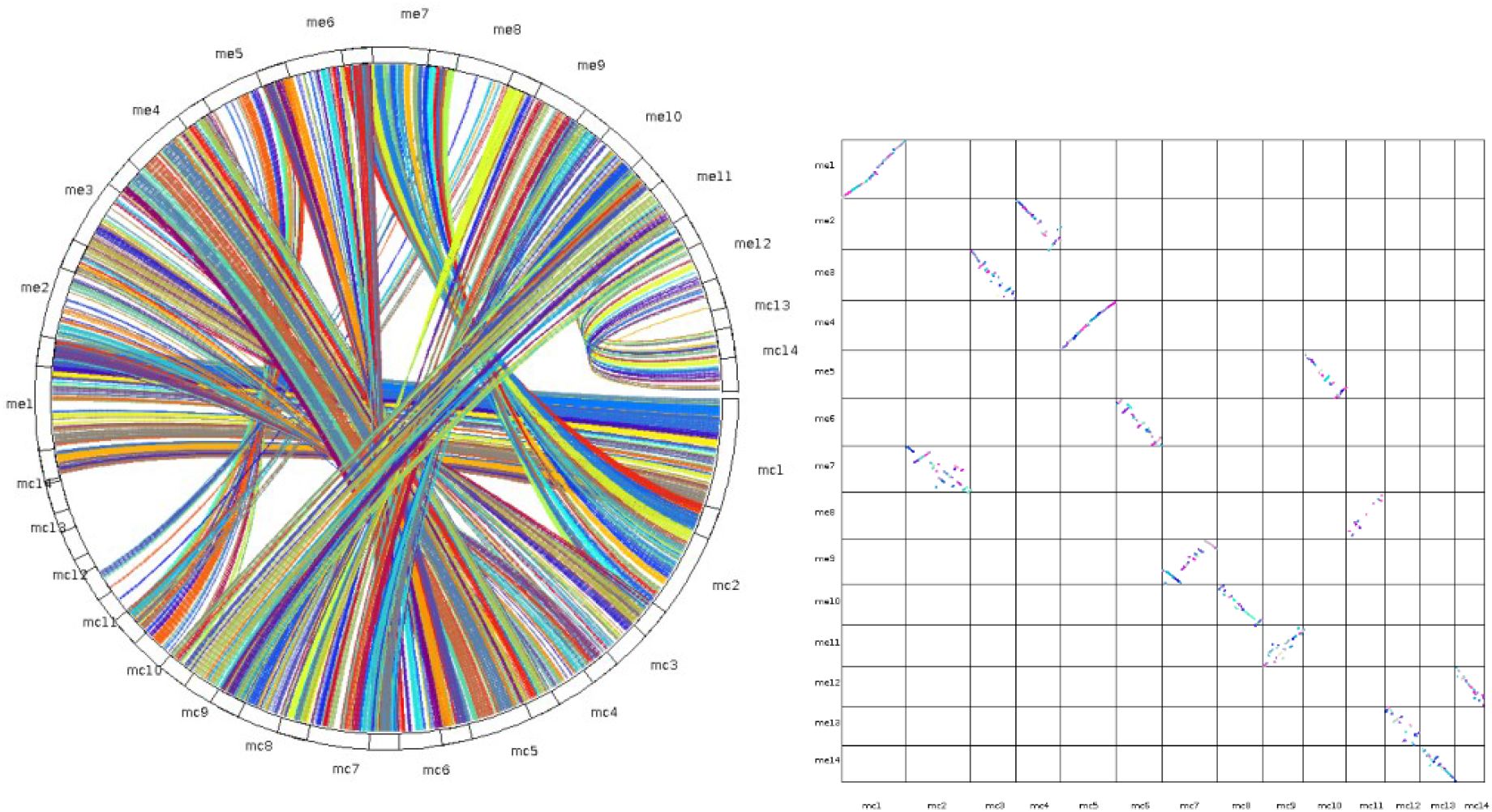
Synteny between the *M. edulis* and *M. coruscus* putative chromosomes.

**Figure 4.**
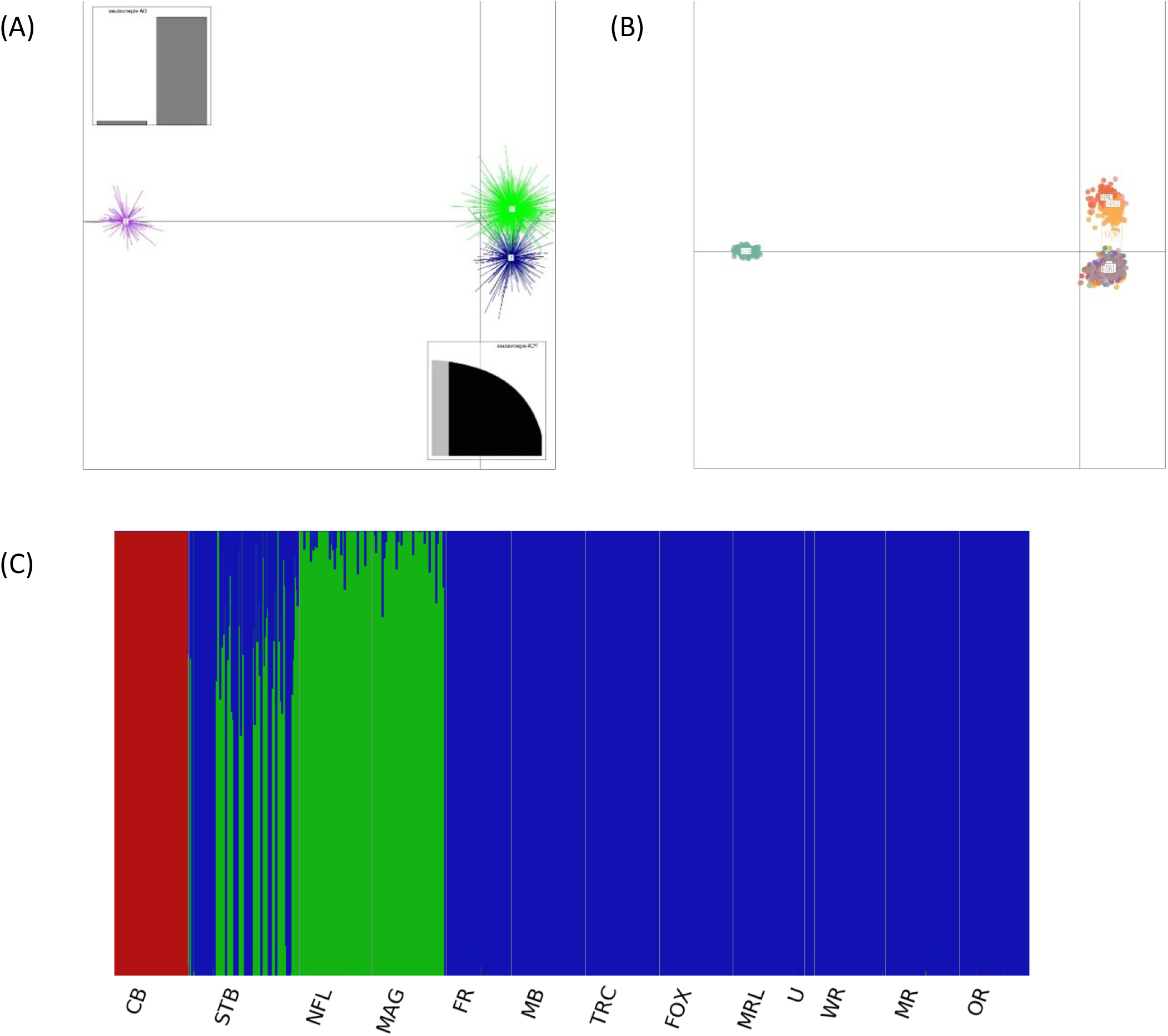
Population structure in the North American Atlantic. (A) DPCA with 6,000 SNPs, (B) PCA with 6,000 SNPs, (C) fastStructure with 6,000 SNPs. Samples from Cape Breton NS (CB), FR, TRC, MB, FOX, MRL, U, WR, MR, OR (PEI), Magdalene Islands QC (MAG), Notre Dame Bay NL (NFL)

The taxonomic status of the “species” in the *Mytilus* species complex remains in debate. Chromosome-level assemblies allow the study of macroevolution of the genome by looking at synteny across species. Herein, we present the synteny analysis between the 14 putative chromosomes of *M. edulis* and *M. corruscus* (Yang et al. 2021) to exemplify how chromosome-level assemblies may allow us to better understand the phylogenetic relationships within the genus *Mytilus*. This analysis shows that some of putative orthologous chromosomes of the 2 species maintain high-levels of collinearity (e.g. chromosomes 1 and 4 from *M. edulis* with LG1 and LG5 from *M. coruscus* respectively) while others present significant levels of re-arrangements (e.g. chromosomes 7 and 11 from *M. edulis* with LG2 and LG9 from *M. coruscus* respectively). An in-depth analysis of chromosome synteny will shed light on the level of collinearity between multiple members of the genus *Mytilus*. Homology between chromosomes is a key element of the viability of hybrids. Reproductive isolation tends to increase during speciation, and these resources will permit further studies on the reproductive compatibility of the species in genus *Mytilus* at the chromosome level.

Here, we present a highly contiguous chromosome assembly for *Mytilus edulis* confirming species-level individual purity through resequencing. To date, our resource has been applied in multiple studies analysing *Mytilus* genome assemblies (Paggeot et al. 2022b; Gallardo-Escarate et al. 2023) and cross-species gene orthology analyses (Saco et al. 2023). The gene annotations produced in this study were generated using Augustus gene model predictions integrating full transcript Isoseq data and applying stringent filtering parameters. This comprehensive approach provides a robust foundation for future cross-species analyses and biological studies on gene function within the *Mytilus* species complex.

## Data Availability

The resulting draft is deposited on NCBI assembly under accession number GCA_019925275.1.

## Funding Statement

“Breeding Better Mussels (Mytilus edulis): developing genomic tools for the implementation of a modern and sustainable mussel Breeding Program” (Grant ID GA-PE-RP3, Genome Canada) The authors would like to acknowledge funding from BBSRC Institute Strategic Programme grants (BBS/E/RL/230001A, BBS/E/RL/230001B and BBS/E/RL/230002B).

## Acknowledgements

Industry Partners: Mussel King (Scott Dockendorff), PEI Aqua Farms (John Paquet), Atlantic Aqua Farms (Terry Ennis).

PEIMSO: Angela MacDonald, Neil McNair, Aaron Ramsey, Jessica Fry

Genome Atlantic: Kristin Tweel, Charmaine Gaudet, Nil d’Entremont

Service Providers: LGC Berlin and UK, Genome Quebec, Dovetail Genomics

## Notes

https://www.ncbi.nlm.nih.gov/data-hub/genome/GCA_019925275.1/

### Competing Interest Statement

The authors have declared no competing interest.

### Summary of Updates

This version was updated with Isoseq data to improve gene model prediction.

